# Assessment of a small-scale fishery: Lane Snapper (*Lutjanus synagris*) using a length metric method

**DOI:** 10.1101/2020.05.13.093856

**Authors:** Liliana Sierra Castillo, Masami Fujiwara

**Author notes:** Author responsible for correspondence: Liliana Sierra^1^, Texas A&M University, Department of Wildlife and Fisheries Sciences.

## Abstract

Small-scale fisheries are hard to assess because of the limited availability of data. Therefore, a method requiring easy-to-obtain catch-data is important for the assessment and management of small-scale fisheries. The objectives of this study were to assess the effect of fishing gear selectivity on a length-based metric method proposed by Froese by estimating three indicators using catch-data from Lane Snapper (*Lutjanus synagris*) collected in Honduras. These indicators are (1) the percentage of mature individuals in the catch, (2) the percentage of fish within the range of estimated optimal lengths to be captured and (3) the percentage of fish larger than the optimal length. These indicators determine the level of overfishing. The indicators were estimated separately for catch-data corresponding to gillnets, and each indicator was estimated with and without selectivity correction. Selectivity and mesh sizes of the fishing gear had a major impact in the estimation of indicators 1 and 2. As for indicator 3, it consistently showed a high level of exploitation. The three estimated indicators suggested that the Lane Snapper fishery in Honduras, is experiencing overfishing. Overall, the method proposed by Froese appears to be promising for the assessment of small-scale fisheries, but it should be used cautiously.

## 1. INTRODUCTION

The status of small-scale fisheries around the world is uncertain because of a lack of adequate data (Ault et al., 2008; Babcock et al., 2013; Babcock et al., 2018; Worm et al., 2009). These fisheries are often not assessed or are assessed inadequately (Aschenbrenner et al., 2017; Carruthers et al., 2014; Froese, 2000; Froese et al., 2012; Levin et al., 2006). Because the stock-assessment models used for the assessment of fisheries are designed for large-scale fisheries, requiring a large amount of high-quality data (Cope & Punt, 2009), it is often very difficult to assess small-scale fisheries using these models (Babcock et al., 2013; Babcock et al., 2018; Froese et al., 2012). Nevertheless, small-scale fisheries are very important globally and still need to be assessed (Andrew et al., 2007; Ray Hilborn et al., 2003; Pauly, 1997).

In order to assess the status of small-scale fisheries, models used to assess large-scale fisheries (such as yield per recruit) are often applied; however, because of the complexity inherent in the models and the data needed to use them, the results are often not satisfactory (Froese, 2004). Thus, it has been suggested that the assessment methods for small-scale fisheries should depend on a simple method using the data that are easy to obtain, such as size/age frequency data (Freitas et al., 2014; Orensanz et al., 2005; Punt et al., 2001; Worm et al., 2009). The analysis of length frequency data can provide important information about shifts in the population age/size structure, which can be indicative of overexploitation (Babcock et al., 2018).

The length metric method proposed by Froese (2004) is one method that is simple and depend on commonly available length frequency data. The method utilizes length data from catches to estimate three indicators of overfishing: (1) the percentage of mature fish present in the length frequency data, (2) the percentage of fish caught within an optimal length, and (3) the percentage of “mega spawners” present in the length frequency data. These indicators are designed to examine the size distribution of fish in the ocean. The logic behind them is that (1) there should be a high percentage of mature individuals (individuals that are above the length where at least 50% of the population has reproduced at least once) in the ocean to avoid recruitment overfishing; (2) there should be a high percentage of individuals that are considered to be within the range of optimal lengths, which are the lengths at which the highest yield can occur; and (3) the percentage of “mega spawners” present in the ocean should be between 20% and 40% in order to conserve large and mature individuals, and thereby avoid growth overfishing (Aschenbrenner et al., 2017; Cope & Punt, 2009; Froese, 2004). Its simplicity makes the method very attractive to fishery managers (Aschenbrenner et al., 2017; Busilacchi et al., 2012; Cope & Punt, 2009; Cury & Christensen, 2005; Francis et al., 2007; Lewin et al., 2006).

These indicators provide important information for fishery assessment (Babcock et al., 2018). Overfishing occurs when the catches in a given period are at maximum sustainable yield level or exceed the catches that surpass desired levels, such as maximum yield-per-recruit (Hilborn & Hilborn, 2011). Overfishing may be categorized into recruitment overfishing, which occurs when recruitment is reduced, or growth overfishing, which happens when too many small fish are caught. Indicators 1 and 3 determine these two types of overfishing that might be occurring in the fishery (Sissenwine & Shepherd, 1986). Indicator 2 helps to achieve the maximum yield of the fishery (Froese, 2004).

Even though using these indicators might be promising for the assessment of small-scale fisheries, one major drawback is the underlying assumption that the catch length composition is representative of the fish in the ocean (Babcock et al., 2013; Cope & Punt, 2009). The data in small-scale fisheries are usually fishery dependent. This is particularly problematic when the catches do not represent the natural populations because of gear selectivity. Gill nets are one of the most size-selective fishing gear (Doll et al., 2014; Gulland, 1987). Thus, the catches may not represent the actual structure of the population of the stock when gill nets are used, which are commonly used in smallscale fisheries (Acosta, 1994; Hamley, 1975; Lobyrev & Hoffman, 2018; McClanahan & Mangi, 2004; Reis & Pawson, 1992; Shoup & Ryswyk, 2016).

Developing an approach for assessing and managing small-scale fisheries with simple methods using readily available data is critically important. The objectives of this study were to determine if the assumption behind the method proposed by Froese (2004) is appropriate for the Lane Snapper (*Lutjanus synagris*) from a small-scale fishery in Honduras and to determine the status of the population of the Lane Snapper (*Lutjanus synagris*) under current management practices. We will use length data collected in Tela, Honduras, as a case study and discuss the adequacy of the length-based method for the assessment of small-scale fisheries.

## 2. METHODS

Sampling was conducted in Tela, Honduras (Figure 1), by the Coral Reef Alliance, the World Wildlife Fund (WWF), and the National University of Honduras (UNAH) as part of a project from the Coral Reef Alliance in coordination with local Honduran authorities, which purpose was to strengthen the capacities for fishermen and the collection of fish data for posterior assessment. For this project, no ethical permissions were required because the surveys and related were done in coordination with local Honduran authorities who manage the areas in Tela. Tela is a coastal town located on the northern coast of Honduras. Data was collected by surveying fishers in the villages of Tornabe, Triunfo de la Cruz, Tela Town, and Miami from 2015 to 2017. The surveys in each village were done until there were no more fishers available to survey. The total number of surveys depended on the number of fishers available and thus was not consistent among days and towns. These towns were selected over others because of their greater number of fishers and accessibility.

**Figure 1.**
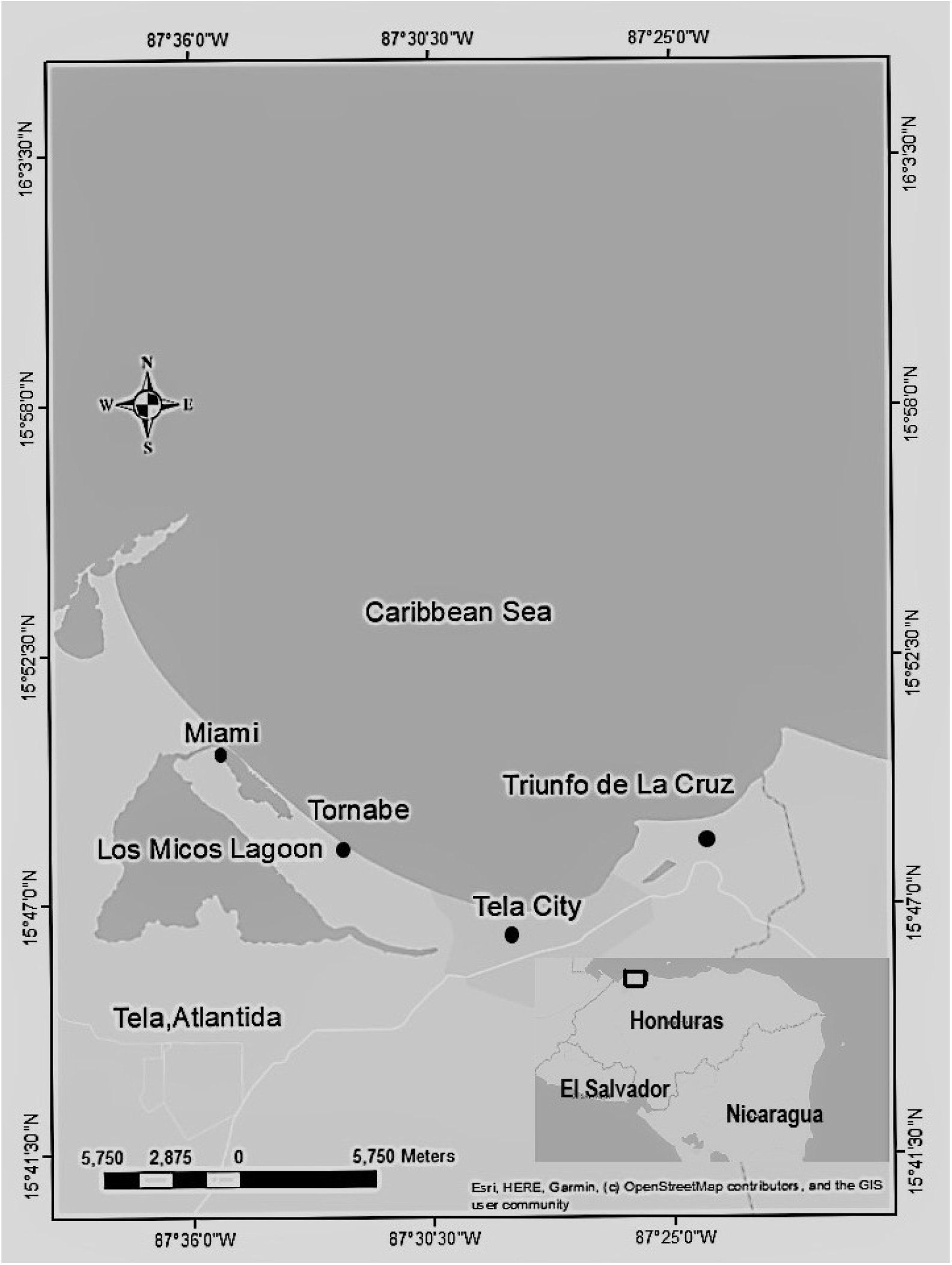
Map of Tela on the northern coast of Honduras and the villages surveyed: Miami, Tornabe, Tela Town, and Triunfo de La Cruz.

In each survey, species, fork length (centimeters) and weight (grams) for each fish, fishing trip duration (minutes), and fishing gear characteristics (gear type and, in the case of gill nets, the specific mesh size) were recorded. The data included many species caught with multiple types of fishing gear. For this study, we analyzed data for Lane Snapper (*Lutjanus synagris*) caught with gill nets of 2” and 3” mesh sizes.

For this study, the length-based method proposed by Froese (2004) was applied to assess the Lane Snapper (*Lutjanus synagris*) fishery in Tela, Honduras. The method consists of the estimations of three indicators that provide the information needed for the assessment of the fishery. The main assumption behind this method is that the length composition of the catch is representative of the length composition of the fish in the ocean.

Each indicator was determined for the data belonging to Lane Snapper (*Lutjanus synagris*) caught with gill nets of 2” and 3” mesh size. The selectivity curve estimated for the same population by Castillo et al. (2018) was used to correct for gear selectivity. The different indicators were described as follows:

### 2.1 Percentage of mature fish present in the length frequency data

The catches of healthy fisheries are expected to include a high percentage of mature individuals (Aschenbrenner et al., 2017; Froese, 2004). For this indicator, we determined the amount and percentage of individuals that were above the *L*_50_, which was the length where 50% of the individuals were able to reproduce. For this study, the average of the different values for *L*_50_ reported in the study in Roatan, Honduras, was used (Berthou et al., 2001).

### 2.2 Percentage of fish caught within an optimal length

The optimal length is the length of individuals caught where the maximum yield is achieved (Froese, 2004), and it was estimated using the expression given by Beverton (1992):

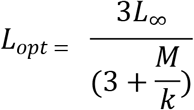

where *L*_∞_ is the maximum length that fish in the population would reach if they were to grow indefinitely, *k* is the growth parameter, and *M* is the natural mortality. For this study, *L*_∞_ of 51.6 cm and *k* of 0.23 were used, taking the values for Lane Snapper (*Lutjanus synagris*) in Puerto Rico (Appeldoorn & Acosta, 1992). Different natural mortalities estimated for Lane Snapper (*Lutjanus synagris*) by Castillo et al. (2018) were used to assess the impact on the optimal length estimation. A range ±10% of the optimal length—as proposed by Babcock et al., (2013), Cope and Punt (2009), and Froese (2004)—was used to determine the percentage of the catch that falls within the optimal length.

### 2.3 Percentage of fish caught that are “mega spawners”

The “mega spawners” in the catch are those fish that are old and are larger than the optimal length (estimated in indicator 2). These fish are of major importance to the fishery because the larger fish tend to be more fecund (Froese, 2004; Gwinn et al., 2015). We estimated the percentage of fish that were larger than the optimal length plus 10%, and considered it to be the percentage of “mega spawners” present in the catch (Froese, 2004). The expected percentages of “mega spawners” should be between 30% and 40% for a healthy stock without overfishing. A percentage of “mega spawners” lower than 20% indicates potential overfishing (Babcock et al., 2018; Froese, 2004).

## 3. RESULTS

### 3.1 Indicator 1: Percentage of mature fish in the length distribution of the catches

Figures 2 and 3 demonstrate the length frequency distribution of the different catches with 2” and 3” mesh size gill nets without selectivity correction. For mesh size 2” (Figure 2), the percentage of mature individuals that are being caught is 13% (when using an *L*_50_ of 24 centimeters), which results in 87% of immature fish in the catch with this mesh size. On the other hand, for mesh size 3” (Figure 3), the percentage of mature individuals reported in the catches is 70% (30% of immature individuals). For this indicator, the percentage of mature individuals should be high (Froese, 2004). As seen in Figures 2 and 3, gill nets of mesh size 3” catch the percentage of mature individuals close to what is recommended, whereas gill nets of mesh size 2” catch a large amount of immature fish.

**Figure 2.**
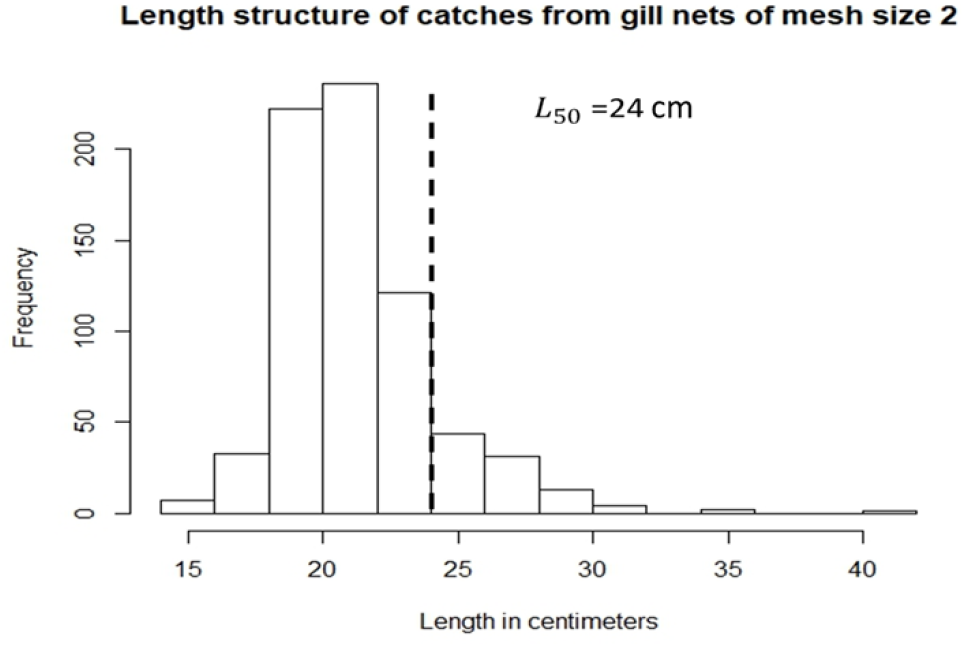
Length distribution of Lane Snapper (*Lutjanus synagris*) caught with gill nets of mesh size 2”; dashed line demonstrates the *L*_50_ used and the frequency of fish that are above and below this.

**Figure 3.**
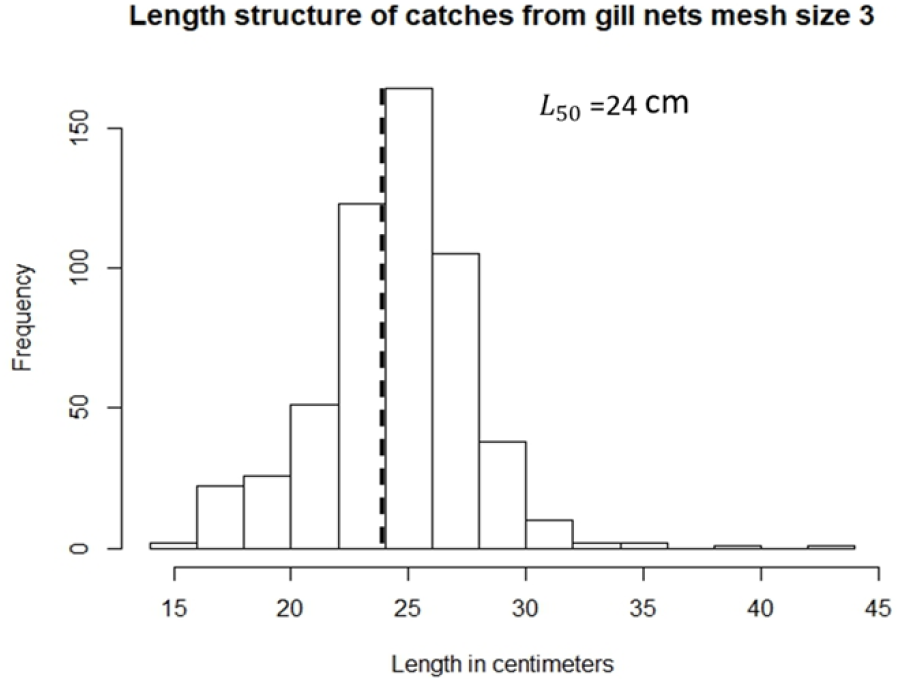
Length distribution of Lane Snapper (*Lutjanus synagris*) caught with gill nets of mesh size 3”; dashed line demonstrates the *L*_50_ used and the frequency of fish that are above and below this.

After correcting for selectivity (i.e., for 2” and 3” mesh sizes) and obtaining the percentage of individuals that are above *L*_50_ (24 cm), there are around 21% of mature individuals in the stock in the ocean.

### 3.2 Indicator 2: Percentage of fish caught at an optimum length

Figures 4 and 5 show the length frequency for each mesh size and the range of lengths considered to be optimum for capture. The different natural mortalities used had an impact on these ranges and thus on the percentages of individuals that were captured at an optimum length (Table 1). As with indicator 1, gill nets of mesh size 2” have low percentages of capturing fish that are in optimum length (these captures range from 3% to 14%. On the other hand, gill nets of mesh size 3” have higher percentages of individuals that are being captured in the optimum lengths, ranging from 10% to 56%.

**Figure 4.**
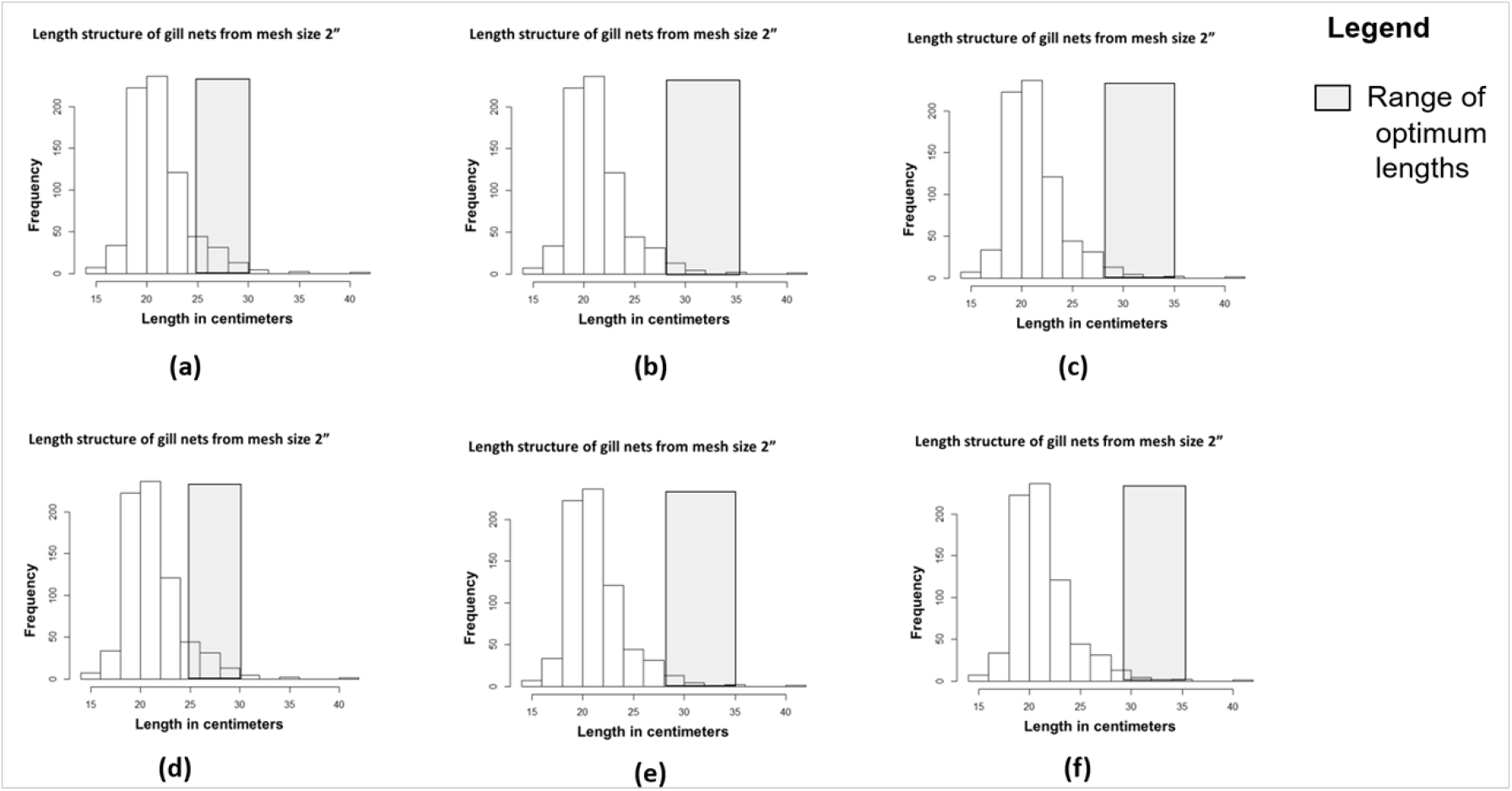
Length frequency distributions of fish caught with mesh size 2” and the optimum range of lengths.

**Figure 5.**
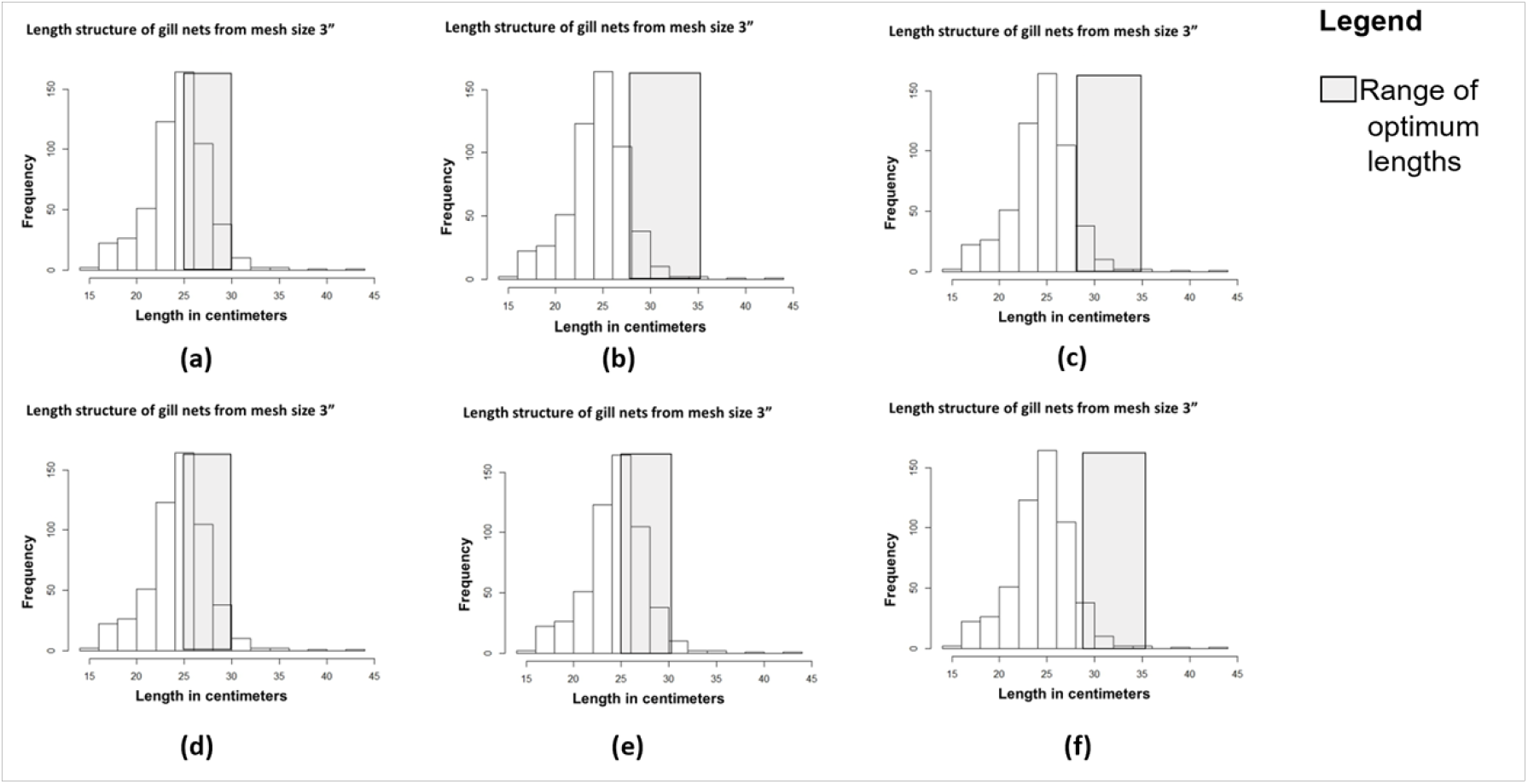
Length frequency distributions for mesh size 3” and the optimum range of lengths.

**Table 1.** Percentage of individuals caught at optimum length with different mesh sizes under different values of assumed natural mortality, with and without selectivity correction.

| **Natural mortality value used** | **Mesh size** | **Percentage of individuals caught at optimum length with selectivity correction** | **Percentage of individuals caught at optimum length without selectivity correction** |
| --- | --- | --- | --- |
| 0.6 | 2" | 20.05% | 12.46% |
| 0.61 | 2" | 20.05% | 12.46% |
| 0.438 | 2" | 11.42% | 4.62% |
| 0.575 | 2" | 21.10% | 12.61% |
| 0.46 | 2" | 11.42% | 4.48% |
| 0.4113 | 2" | 7.34% | 2.66% |
| 0.6 | 3" | 20.41% | 56.12% |
| 0.61 | 3" | 20.41% | 56.12% |
| 0.438 | 3" | 5.25% | 17.73% |
| 0.575 | 3" | 20.80% | 57.40% |
| 0.46 | 3" | 5.20% | 17.55% |
| 0.4113 | 3" | 2.77% | 9.51% |

When correcting for the selectivity that gill nets have, the results indicate that approximately 20% of fish in the ocean would be at the optimum length to be caught (Table 1).

### 3.3 Indicator 3: Percentage of fish caught that are considered to be “mega spawners”

The percentages of mega spawners that are being caught with both mesh sizes (2” and 3”) were very low (0 and 0.18, respectively). For this indicator, the percentage of mega spawners in the catch should be greater than 20% for a stock to be considered healthy (Froese, 2004). Similarly, when taking the selectivity into consideration, there are about 5% of mega spawners in the population in the ocean.

## 4. DISCUSSION

The method proposed by Froese (2004) to assess a fish stock may not be adequate for the assessment of small-scale fisheries without correction for the size-selectivity of the fishing gear. The method assesses the size distribution of the stock in the ocean based on size distribution in the catch. Therefore, the assumption behind this method is that there is no gear selectivity affecting the catches of the fishermen, and thus the catches are representative of the fish stock (Babcock et al., 2018; Cope & Punt, 2009). However, most small-scale fisheries use size-selective fishing gear, and this was the case with our data from the small-scale fishery in Honduras.

When we used the catches from fishing gear that targets smaller fish (e.g., Lane Snapper data collected with 2” mesh size gill nets from Honduras) to estimate the indicators without consideration of the impact of the selectivity, we will obtain lower values for indicators 1 and 2 (Table 2). Using these indicators without selectivity correction to make management decisions can be problematic (Cope & Punt, 2009). For example, indicator 1 without selectivity correction suggests that only 13% of the stock is mature (using the catch as a representation of the stock); however, the same indicator after correcting for gear selectivity suggests that 21% of the stock is mature.

**Table 2.** Estimation of indicators with and without selectivity correction using catches from mesh size 2” gill nets and 3” gill nets.

| Size of mesh size gill nets | Indicator | Estimation of indicators not correcting for selectivity | Estimation of indicators correcting for selectivity |
| --- | --- | --- | --- |
| 2” | Indicator 1 | 18.60% | 21.85% |
| 2” | Indicator 2 | 12.46% | 20.05% |
| 2” | Indicator 2 | 4.62% | 11.42% |
| 2” | Indicator 2 | 12.61% | 21.10% |
| 2” | Indicator 2 | 4.48% | 11.42% |
| 2” | Indicator 2 | 2.66% | 7.34% |
| 2” | Indicator 3 | ~0 | ~5% |
| 3” | Indicator 1 | 70% | 21% |
| 3” | Indicator 2 | 56.12% | 20.05% |
| 3” | Indicator 2 | 56.12% | 20.05% |
| 3” | Indicator 2 | 17.73% | 11.42% |
| 3” | Indicator 2 | 57.40% | 21.10% |
| 3” | Indicator 2 | 17.55% | 11.42% |
| 3” | Indicator 2 | 9.51% | 7.34% |
| 3” | Indicator 3 | 0.18% | ~5% |

On the contrary, with fishing gear that target larger sizes of fish (e.g., Lane Snapper data collected with 3” mesh size gill nets from Honduras), the catch—and thus length—structures consist of larger age classes. As a result, when selectivity is not taken into consideration, the estimation of indicators 1 and 2 will be higher (Table 2). Indicator 1 (without selectivity correction) suggests that 70% of the individuals in our catch are mature; this may suggest that the stock is healthy. However, the percentage of mature individuals, when taking selectivity into account, is 21%, and thus recommending the use of mesh size 3” (or larger) might affect the older age classes of the stock. This will cause problems when interpreting the indicators to make management recommendations (Babcock et al., 2013; Babcock et al., 2018; Cope & Punt, 2009).

Although indicator 3 values for both mesh sizes (2” and 3”) are lower when compared with the indicator estimated after correction for gear selectivity (Table 2), the estimation can also be higher if gear with much larger mesh size is used. It can potentially suggest that the stock may be healthy, but it does not reflect the true condition of the stock (Babcock et al., 2018).

The status of the Lane Snapper in Tela, Honduras, based on the three indicators after correcting for the size-selectivity of the gill nets, is an overfished stock. With indicator 1, the percentage is 21%, which suggests that a small percentage of fish are mature. Additionally, indicator 3 shows that the percentage of mega spawners is currently 5%, when it should be greater than 30%. Therefore, recruitment overfishing and growth overfishing are occurring with the stock (Froese, 2004; Froese et al., 2018; Froese et al., 2012). Agreeing with Castillo et al. (2018), there seems to be high fishing pressure (resulting in high fishing mortalities), which is currently causing the overfishing. Because of this, we recommend that the managers eliminate the use of mesh size 2” gill nets entirely. Even though they are illegal to use in Honduras (as stated by law), there are many fishermen still using them. It may be difficult to enforce the ban of mesh size 2” gill nets in Honduras because of the unique sociological and economic characteristics surrounding the fishery, but we believe it is possible—by working side by side with the fishers and local organizations—to implement this recommendation (Carbajal et al., 2017)

At the same time, the fishing pressure on the older age classes should be reduced. To achieve this goal, we recommend closing the areas where older age classes of Lane Snapper spend most of their time and protecting these areas from gill net fisheries. This can be implemented as part of the regulations of the marine protected area (MPA) recently established in the area. In addition, the overall fishing effort should be decreased by reducing the number of gill nets the fishers can use in each fishing trip.

In general, when the method proposed by Froese (2004) is used, the results should be interpreted with caution. Without consideration of the selectivity of fishing gear, the indicators may be overestimated or underestimated. However, we also understand that the simplicity of the method, which is the major advantage of it, is undermined by including an additional step in calculating the indicators taking selectivity into consideration. Furthermore, there may not be adequate data to correct for gear selectivity; however, it is very important to correct for it. When it is necessary to use the indicators estimated without correcting for gear selectivity, it is important to know that indicators will have higher values when the fishing gear selects smaller fish and lower values when the gear selects larger fish. Thus, the management decision should be adjusted accordingly. It may be necessary to take a precautionary approach when the fishing gear is highly selective to certain sizes in order to protect a stock. Overall, the length metric method proposed by Froese (2004) might seem promising for small-scale fisheries because the data that is needed can be easily obtained, and it provides useful information about the stock.

## 5. ACKNOWLEDGEMENTS

We would like to acknowledge the Coral Reef Alliance and the World Wildlife Fund (WWF) for providing the funding that made this study possible. Also, we would like to acknowledge the National University of Honduras (UNAH) for making the collection of data and preliminary analysis of data possible. Finally, without the help of all the fishers and their families from the different communities, this study could not have been realized. MF was supported, in part, by NSF OCE 1656923. The map in this article was created using ArcGIS^®^ software by Esri. ArcGIS^®^ and ArcMap™ are the intellectual property of Esri and are used herein under license. Copyright © Esri. All rights reserved. For more information about Esri^®^ software, please visit www.esri.com.

